# Coordination difficulties, IQ and psychopathology in children with high-risk Copy Number Variants

**DOI:** 10.1101/662833

**Authors:** Adam C Cunningham, Jeremy Hall, Michael J Owen, Marianne B M van den Bree

## Abstract

**Background:** The prevalence and impact of motor coordination difficulties in children with Copy Number Variants that are associated with high risk of neurodevelopmental disorder (ND-CNVS) remain unknown. The present study aims to advance understanding of motor coordination difficulties in children with ND-CNVs and establish relationships with IQ and psychopathology.

**Methods:** 169 children with a ND-CNV (67% male, median age 8.88 years, range 6.02-14.81) and 57 closest-in-age unaffected siblings (controls; 55% male, median age 10.41 years, SD=3.04, range 4.89-14.75) were assessed with the Developmental Coordination Disorder Questionnaire, alongside psychiatric interviews, and standardised assessments of IQ.

**Results:** 91% of children with an ND-CNV screened positive for coordination problems, compared to 19% of unaffected sibling controls (OR=42.53, p<.001). There was no difference in coordination ability between ND-CNV genotypes (F=1.47, p=.184). Poorer motor coordination in the ND-CNV group was associated with greater numbers of ADHD (p=.021) and autism spectrum disorder trait (p<.001) symptoms, along with lower full-scale (p=.011), performance (p=.015), and verbal IQ (p=.036). Mediation analysis indicated that coordination ability was a full mediator of anxiety symptoms (69% mediated, p=.012), and a partial mediator of ADHD (51%, p=.001) and ASD trait symptoms (66%, p<.001) along with FSIQ (40%, p=.002) PIQ (40%, p=.005) and VIQ (38%, p=.006) scores.

**Conclusions:** The findings indicate that poor motor coordination is highly prevalent and closely linked to risk of mental health disorder and lower intellectual function. Future research should explore whether early interventions for poor coordination ability could ameliorate neurodevelopmental risk more generally.

## Background

Difficulties with motor skills can have serious consequences for a child’s independence and daily functioning (Van der Linde et al., 2015) and there is evidence that these negative effects can persist into adulthood (Kirby, Sugden, & Purcell, 2014; Kirby, Williams, Thomas, & Hill, 2013).

Difficulties with co-ordinated movement are often seen in combination with other neurodevelopmental disorders such as ADHD and autism spectrum disorder (ASD). For example, it has been estimated that up to 50% of children with Developmental Coordination Disorder (DCD), a neurodevelopmental disorder characterised by functional difficulties with coordinated movement, also possess a diagnosis of ADHD, usually of the inattentive subtype (Kadesjö & Gillberg, 1999; Kaiser, Schoemaker, Albaret, & Geuze, 2015). Individuals with DCD also often show problems in neurocognition, particularly in executive functioning (Wilson, Ruddock, Smits-Engelsman, Polatajko, & Blank, 2013).

A range of genomic disorders such as those caused by sub-microscopic deletions or duplications of chromosomal regions including 1q21.1, 16p11.2 or 22q11.2 have been associated with the development of conditions such as ADHD, ASD, schizophrenia and intellectual disability (Torres, Barbosa, & Maciel, 2015). These chromosomal abnormalities are termed Copy Number Variants (CNVs), as they change the number of copies of genes contained on the affected area of the chromosome. While there is strong evidence that many CNVs are associated with a high risk of developing neurodevelopmental disorder (ND), including psychopathology (referred to hereafter as ND-CNVs), penetrance is often incomplete and variable. This means that while some individuals will display many complex symptoms, others may show few or none (Crawford et al., 2018).

Previous research by our group has found that approximately 80% of children with 22q11.2 Deletion Syndrome (22q11.2DS) show poor coordination of movement (Cunningham et al., 2017). Our findings also indicated that the children who showed poorer motor coordination had higher risks of ADHD, ASD and anxiety symptoms and lower mean IQ. However, there is very little research into coordination difficulties in children with other ND-CNVs. It is therefore unclear if individuals with other ND-CNVs experience similar coordination difficulties, or if certain ND-CNVs confer greater risk for coordination difficulties than others. Similarly, it is unknown if the links between coordination difficulties and other neurodevelopmental symptoms that we found for 22q11.2DS are also present on other high-risk ND-CNVs. More generally, the links between coordination difficulties and other neurodevelopmental problems are not well understood.

Finally, it is not known to what extent motor coordination disorder mediates the effects of carrying a ND-CNV on subsequent neurodevelopmental impairment. This idea is supported by theories that the early development of motor skills influences the development of other higher cognitive processes (Wilson, 2002). Motor skills develop very early in life and it follows that difficulties with interacting with and exploring one’s environment impact on the development of other skills. For example, it has been suggested that poor motor development will impact on skills such as the representation of abstract concepts (Piaget, 1954), mathematics (Giles et al., 2018), and language ability (Rowe, Özçalışkan, & Goldin-Meadow, 2008).

With these ideas in mind, we assessed motor coordination, IQ and psychopathology in a large group of children with CNVs associated with the development of neurodevelopmental disorder in order to test the following hypotheses: 1) Children with ND-CNVs have an increased rate of motor coordination difficulties compared to unaffected sibling controls. We base this hypothesis on research where neurodevelopmental problems have been found to be associated with increased risk of motor coordination difficulties in non-genotyped samples (Kadesjö & Gillberg, 1999; Kaiser et al., 2015; Pratt & Hill, 2011; Skirbekk, Hansen, Oerbeck, Wentzel-Larsen, & Kristensen, 2012; Sumner, Leonard, & Hill, 2016), as well as evidence from 22q11.2DS (Cunningham et al., 2017); 2) Motor coordination ability will differ across genotypes, as different ND-CNV’s affect different genes in different areas of the genome; 3) Poor coordination will be related to increased levels of psychopathology and lower IQ in children with ND-CNVs, similar to the pattern seen in non-CNV populations (Harrowell, Hollén, Lingam, & Emond, 2017; P. H. Wilson et al., 2013) as well as 22q11.2DS (Cunningham et al., 2017) and; 4) The risk of psychopathology and low IQ posed by carrying a ND-CNV is partially indirect, via motor coordination ability, or in other words, motor coordination ability will mediate the relationship between ND-CNV status and psychopathology and IQ. This hypothesis is supported by findings that appropriate development of motor skills is required for the development of higher-order cognitive skills (Giles et al., 2018; Rowe et al., 2008; Wilson, 2002).

## Methods

### Participants

One-hundred and sixty-nine participants with a range of ND-CNVs took part (67% male, median age: 8.88 years, range: 6.02-14.81, see Supplementary Table 1 for CNVs included), as well as 57 closest-in-age unaffected siblings (controls; 54% male, median age: 1.41 years, range: 5.89-14.75). Families were recruited through UK Medical Genetics clinics as well as word of mouth and the charities Unique and MaxAppeal!. Informed and written consent was obtained prior to recruitment from the carers of the children and recruitment was carried out in agreement with protocols approved by the London Queens Square NRES Committee. Individual ND-CNV genotypes were established from medical records as well as in-house genotyping at the Cardiff University MRC Centre for Neuropsychiatric Genetics and Genomics using microarray analysis. Information about medical comorbidities including congenital heart defects, epilepsy, and premature birth, along with medication use.

**Table 1.** Demographic and summary statistics of participants

|  | ND-CNV (N=169) | Control (N=72) | p value |
| --- | --- | --- | --- |
| **Income** |  |  |  |
| N-Miss | 18 | 16 |  |
| <=£19,999 | 59 (39.1%) | 12 (21.4%) |  |
| £20,000 - £39,999 | 49 (32.5%) | 27 (48.2%) |  |
| £40,000 - £59,999 | 22 (14.6%) | 8 (14.3%) |  |
| £60,000 + | 21 (13.9%) | 9 (16.1%) |  |
| **Education** |  |  |  |
| N-Miss | 5 | 10 |  |
| High | 45 (27.4%) | 16 (25.8%) |  |
| Low | 40 (24.4%) | 17 (27.4%) |  |
| Middle | 70 (42.7%) | 28 (45.2%) |  |
| No School Leaving Exams | 9 (5.5%) | 1 (1.6%) |  |
| **Ethnicity** |  |  |  |
| N-Miss | 6 | 10 |  |
| European | 150 (92.0%) | 56 (90.3%) |  |
| Other | 13 (8.0%) | 6 (9.7%) |  |
| **Age** |  |  | < .001<sup>2</sup> |
| N-Miss | 0 | 0 |  |
| Median (Q1, Q3) | 8.877 (7.116, 10.958) | 10.408 (8.801, 12.364) |  |
| Range | 6.016 - 14.810 | 5.894 - 14.753 |  |
| **Gender** |  |  | .062<sup>1</sup> |
| N-Miss | 0 | 0 |  |
| M | 113 (66.9%) | 39 (54.2%) |  |
| F | 56 (33.1%) | 33 (45.8%) |  |
| **Premature** |  |  | .010<sup>1</sup> |
| N-Miss | 11 | 7 |  |
| No | 72 (45.6%) | 42 (64.6%) |  |
| Yes | 86 (54.4%) | 23 (35.4%) |  |
| **CHD** |  |  | .008<sup>1</sup> |
| N-Miss | 6 | 2 |  |
| No | 139 (85.3%) | 68 (97.1%) |  |
| Yes | 24 (14.7%) | 2 (2.9%) |  |
| **Epilepsy** |  |  | < .001<sup>1</sup> |
| N-Miss | 5 | 2 |  |
| No | 138 (84.1%) | 70 (100.0%) |  |
| Yes | 26 (15.9%) | 0 (0.0%) |  |
| **DCDQ Score** |  |  | < .001<sup>2</sup> |
| N-Miss | 0 | 0 |  |
| Median (Q1, Q3) | 32.000 (25.000, 41.000) | 69.500 (58.500, 74.000) |  |
| Range | 15.000 - 73.000 | 19.000 - 75.000 |  |
| **FSIQ** |  |  | < .001<sup>3</sup> |
| N-Miss | 12 | 3 |  |
| Mean (SD) | 81.898 (14.997) | 99.246 (12.838) |  |
| Range | 51.000 - 128.000 | 70.000 - 139.000 |  |
| **PIQ** |  |  | < .001<sup>3</sup> |
| N-Miss | 12 | 3 |  |
| Mean (SD) | 86.146 (15.276) | 101.681 (13.376) |  |
| Range | 54.000 - 132.000 | 75.000 - 131.000 |  |
| **VIQ** |  |  | < .001<sup>3</sup> |
| N-Miss | 11 | 3 |  |
| Mean (SD) | 80.835 (15.306) | 97.130 (14.214) |  |
| Range | 55.000 - 123.000 | 63.000 - 141.000 |  |
| **ADHD Symptom Count** |  |  | < .001<sup>2</sup> |
| N-Miss | 0 | 0 |  |
| Median (Q1, Q3) | 6.000 (2.000, 11.000) | 0.000 (0.000, 1.000) |  |
| Range | 0.000 - 17.000 | 0.000 - 17.000 |  |
| **ASD Trait Scores** |  |  | < .001<sup>2</sup> |
| N-Miss | 0 | 0 |  |
| Median (Q1, Q3) | 18.000 (11.000, 23.000) | 3.000 (2.000, 6.000) |  |
| Range | 0.000 - 33.000 | 0.000 - 34.000 |  |
| **Anxiety Symptom Counts** |  |  | < .001<sup>2</sup> |
| N-Miss | 0 | 0 |  |
| Median (Q1, Q3) | 2.000 (0.000, 8.000) | 0.000 (0.000, 1.000) |  |
| Range | 0.000 - 25.000 | 0.000 - 17.000 |  |
| **ODD Symptom Count** |  |  | < .001<sup>2</sup> |
| N-Miss | 0 | 0 |  |
| Median (Q1, Q3) | 2.000 (1.000, 4.000) | 0.000 (0.000, 2.000) |  |
| Range | 0.000 - 7.000 | 0.000 - 6.000 |  |

### Functional coordination impairment-The Developmental Coordination Disorder Questionnaire (DCDQ)

The DCDQ (Wilson et al., 2009) was completed by the primary carer. It is designed to screen for functional motor coordination impairments in children 5–15 years old. The DCDQ is widely used and validated (Cunningham et al., 2017; Wilson et al., 2009). In general, lower scores indicate greater coordination difficulties. Items probe fine and gross motor skills. It yields a total score as well as separate scores for three subscales: control during movement, fine motor/handwriting and general coordination. The total score was used as a continuous measure of coordination ability. In addition, participants were categorised into those with poor coordination, (scoring positive on the DCDQ) and those without (scoring negative) using age dependant scoring thresholds.

### IQ assessment

Full scale, verbal and performance IQ (FSIQ, VIQ and PIQ) was obtained by administering the Wechsler Abbreviated Scale of Intelligence (Four subtests: matrix reasoning, block design, vocabulary, similarities) (WASI) (Wechsler, 1999).

### Psychiatric assessment

The social communication questionnaire (SCQ) (Rutter, Bailey, & Lord, 2003a) was used to screen for ASD trait symptoms. A score of 15 or greater is considered suggestive of ASD (Rutter, Bailey, & Lord, 2003b). The SCQ consists of three subscales: repetitive behaviour, social interactions and communication and scores across these can be combined to obtain a total score.

ADHD, anxiety and ODD symptoms were assessed using the semi-structured Child and Adolescent Psychiatric Assessment (CAPA) (Angold et al., 2009). The interview was conducted by trained psychologists with the primary caregiver. Interviews were audiotaped, and DSM-5 diagnosis obtained during consensus meetings led by a child and adolescent psychiatrist. We did not consider diagnoses to be mutually exclusive. Anxiety symptoms included any symptom of generalised anxiety disorder, social phobia, specific phobia, separation anxiety, panic disorder with and without agoraphobia, agoraphobia and obsessive-compulsive disorder.

### Statistical analysis

In order to investigate the effect of having an ND-CNV on rates of screening positive for coordination problems, we carried out a chi-squared test comparing rates of positive and negative screening between each group.

Additionally, to investigate coordination ability (DCDQ score) we conducted a linear mixed effect model where the continuous DCDQ total score was predicted by ND-CNV status with age as a covariate, and family membership as a random effect, as the children with ND-CNVs and controls were siblings.

As a sensitivity analysis, this mixed effect model was also run with gender, maternal education, family income, presence of congenital heart defects, epilepsy, premature birth and medication use as additional covariates, in order to identify if these variables should be included as covariates in subsequent analyses.

An ANCOVA was used to investigate the extent to which DCDQ total score was explained by ND-CNV genotype. These analyses were based on the ND-CNV groups with 10 or more individuals available (15q11.2 deletion, 15q13.3 deletion, 16p11.2 deletion, 16p11.2 duplication, 1q21.1 deletion, 1q21.1 duplication, 22q11.2 deletion, 22q11.2 duplication and 2p16.3 deletion (NRXN1)). Age was entered as covariates. A similar ANCOVA was used to investigate the effect of deletion or duplication of genetic material across the ND-CNV group, with age.

In order to investigate the relationship between DCDQ score and psychopathology or IQ in children with CNVs, hierarchical regression models were constructed where DCDQ total score was predicted first by age, then the relevant psychopathology or IQ variable at the second step. For the analysis including IQ, we ran models with both standardised IQ (i.e, full scale IQ, verbal IQ and performance IQ scores) as well as raw IQ subtest scores. The latter were included because of the potential for coordination ability to impact on IQ test performance, particularly in the block design task, where scores are assigned based on time taken to complete patterns. Poor for age motor ability may impede performance on this task, as constructing the patterns may take longer, even if the method is correct.

Additionally, mediation analyses (Baron & Kenny, 1986) were run to investigate whether the relationship between ND-CNV status and psychopathology or IQ scores were mediated by motor coordination ability with age as a covariates (Figure 1). The R package “mediation” v4.4.7 (Tingley, Yamamoto, Hirose, Keele, & Imai, 2014) was used to conduct the mediation analyses. This provides point estimates of the average causal mediation effects (ACME), average direct effects (ADE) and total effects (ADE+ACME) and the proportion of the total effects that are accounted for by the indirect path (proportion mediated). ACME corresponds to the indirect effect through the mediator, while the ADE corresponds to the direct effect of the independent variable on the outcome. Confidence intervals and p-values were obtained using nonparametric bootstrapping with 5000 simulations (Imai, Keele, & Tingley, 2010). The results of the analysis can be interpreted in the following ways, if a significant total effect, and direct effect (ADE) but insignificant indirect effect (ACME) are found, the mediator has no effect on the outcome. If a significant total effect, significant indirect effect, but insignificant direct effect is found, the mediator is fully mediating the effect of the independent variable on the outcome. Finally, if a significant total effect, and significant direct and indirect effects are found, the mediating variable is a partial mediator of the effect of the independent variable on the outcome. Importantly, if the total effect is not significant, there is no evidence that the independent variable has an effect on the outcome.

**Figure 1.**
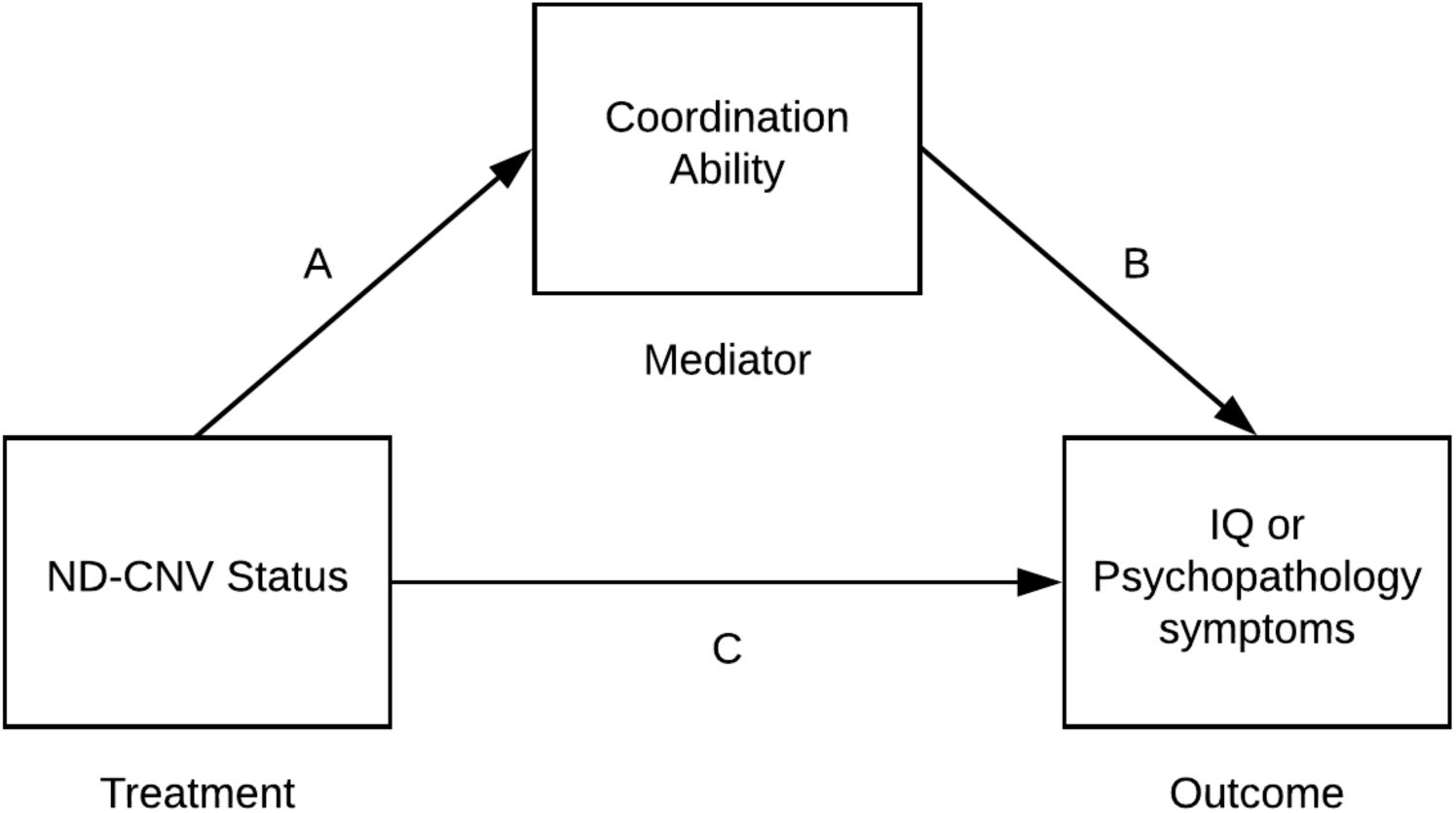
The mediation model we tested to investigate the associations between ND-CNV status (0=sibling control; 1=ND-CNV) and outcome, via a direct (Path C) and indirect pathway (A-B). Pathways A-B estimate to what extent the link between CNV status and outcome can be accounted for by an indirect link via coordination ability (Developmental Coordination Disorder Questionnaire Score) mediator.

To validate that the mediation analyses was robust to the path being investigated, a second set of models investigating if FSIQ mediated the effect of ND-CNV status on psychopathology were also constructed and tested using the same method.

Sensitivity analyses were carried out to ensure that indirect effects were robust to violations of the assumption of sequential ignoreability (effect of unmeasured confounding variables). For these sensitivity analyses, the level of confounding due to unmeasured confounders was represented by the correlation between the residuals (error terms) from the mediator and the outcome models, denoted ρ (rho). If ρ=0 there is no correlation between residuals, and this can be interpreted as no unmeasured confounding. By varying levels of ρ between values of −1 and +1 we can explore how any detected indirect effect is influenced by unmeasured confounders. The key outcome of these sensitivity analyses is the level of ρ for which the indirect effect becomes nonsignificant, giving a measure of how strong the effect of any unmeasured confounding variables would need to be in order to invalidate the estimated indirect effect (ACME).

All analyses were carried out on Mac OSX and R v3.5.3. Number of individuals with information for each assessment may differ, due to not successfully completing assessments.

## Results

Descriptive statistics for the individuals that took part is presented in Table 1. The ND-CNV group was significantly younger than the unaffected sibling controls but had a similar proportion of males and females. Due to this difference in age between the groups, and the fact that DCDQ total score was correlated with age (r=.26, p<.001) in children with an ND-CNV, age was included as a covariate in all analyses.

Eight children with an ND-CNV were receiving sodium valproate for epilepsy, six were receiving methylphenidate, three were taking carbamazepine, three were on risperidone, two atomoxetine, two on levetiracetam, one on clobazam, two on ethosuximide, one on clobazam, one on lamotrigine, one on fluvoxamine, one on guanfacine, one on fluoxetine and one on nitrazepam. No other relevant medication use was noted. Notably, 54% of children with an ND-CNV were born before 37 weeks, 54% had a congenital heart defect, and 16% reported a history of seizures.

### Hypothesis 1: Is there a difference in rates of indicated DCD between CNV carriers and Controls?

Ninety-one percent (154/169) of children with an ND-CNV and 19% (14/72) controls screened positive for coordination problems (OR=42.53, χ^2^=122.86, p=<.001).

Children with an ND-CNV had lower DCDQ total scores than controls (Table 1), and a linear mixed model where DCDQ total score was predicted by ND-CNV status and age with family membership as a random effect, found that ND-CNV status was predictive of DCDQ total score (b=28.98, p<.001), along with age (b=0.78, p=.022) were predictive of DCDQ total score.

Addition of gender, family income, maternal education, or CHD, epilepsy, premature birth or medication use in the mixed effect models as covariates revealed that they had no effect on screening for coordination problems or DCDQ score. Therefore, these were not included as covariates in any subsequent analyses.

Across all ND-CNV genotypes we studied, rates of coordination problems were high, with the lowest rate (77.8%) found in children with 15q13.3 duplication, whilst all (100%) children with 15q11.2 duplication, 16p11.2 distal duplication, 16p11.2 duplication, 1q21.1 deletion, 1q21.1 duplication, 22q11.2 distal deletion, deletion of 9q34.3 (Kleefstra Syndrome), TAR deletion and TAR duplication screened positive (Supplementary Table 1).

### Hypothesis 2: Does coordination score differ by genotype?

After including age, ND-CNV genotype was not a significant predictor of DCDQ score (F=1.47, df=7, p=.184, η^2^=.069) in those ND-CNV groups with 10 individuals or more. These findings are consistent with the null hypothesis that coordination ability is similar regardless of ND-CNV genotype. In addition, type of CNV, (deletion (n= 101) or duplication (n=68)) of material also had no effect on DCDQ score. (F=.67, df=1, p=.413, η^2^=.003)

### Hypothesis 3: Is coordination related to psychopathology and IQ in children with an ND-CNV?

Children with ND-CNV’s displayed higher levels of psychopathology symptoms than siblings (Table 1). When investigating the links between coordination difficulties and psychopathology within the ND-CNV group, we found that worse coordination ability was associated with a greater number of ADHD, and ASD traits, but not anxiety or ODD symptoms. In all models, age was a significant covariate, with older children having better coordination ability (Table 2).

**Table 2.**
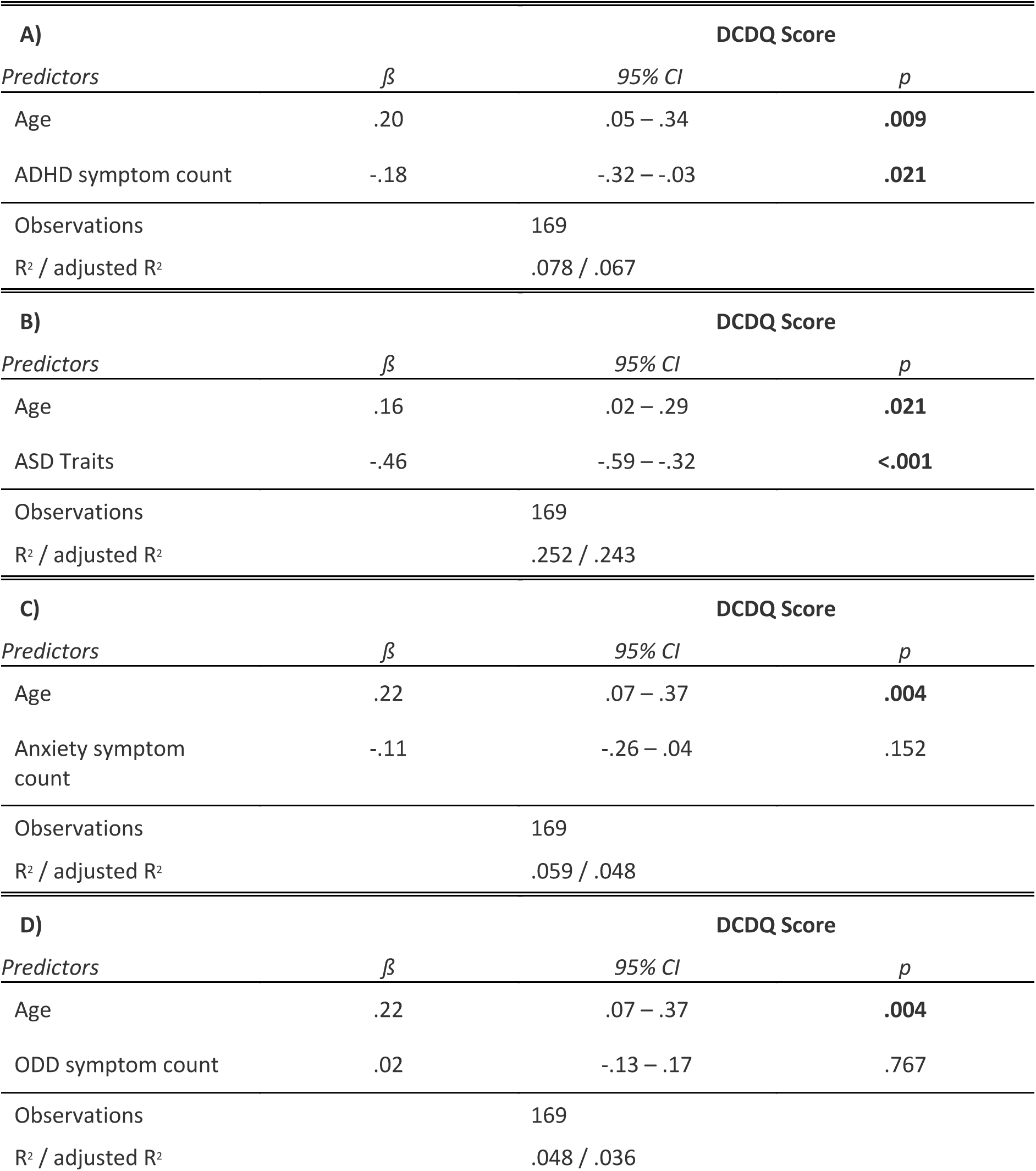
Regression results for DCDQ score predicted by A) ADHD symptom counts, B) ASD traits, C) Anxiety Symptoms, D) Oppositional defiant Disorder (ODD) symptoms, with age as a covariate.

Average FSIQ, PIQ and VIQ were lower in the ND-CNV group (Table 1). Within children with ND-CNV’s, worse coordination was associated with lower FSIQ, PIQ and VIQ, with age as a significant covariate (Supplementary Table 2 A-C). When investigating if DCDQ score was associated with individual raw subtest scores, we found that better matrix reasoning, but not block design, similarities or vocabulary performance was associated with better coordination (Supplementary Table 2 D-G).

### Hypothesis 4: Do coordination difficulties mediate the relationship between ND-CNV group status and psychopathology or IQ?

Mediation analysis indicated that coordination ability (DCDQ score) was a full mediator of the effect of having and ND-CNV on anxiety symptoms (69% proportion mediated) and was a partial mediator of ADHD symptoms (51% proportion mediated) and ASD traits (66% proportion mediated). While no evidence for mediation was found for ODD symptoms (Table 3.).

**Table 3.** Results of mediation analysis on the effect of having an ND-CNV on A) ADHD symptom counts, B) ASD traits, C) Anxiety symptoms, D) ODD symptoms with coordination ability as a mediator. Average Causal Mediation Effects corresponds to the indirect effect through coordination ability, while Average Direct effects corresponds to the direct effect of having an ND-CNV on the psychopathology variable of interest.

| **A) ADHD symptom counts** | **Estimates** | **Lower 95% CI** | **Upper 95% CI** | **p** |
| --- | --- | --- | --- | --- |
| Average Causal Mediation Effects | 2.54 | 1.04 | 4.24 | **.001** |
| Average Direct Effects | 2.39 | .13 | 4.44 | **.042** |
| Total Effects | 4.93 | 3.67 | 6.10 | **<.001** |
| Prop.Mediated | .51 | .20 | .97 | **.001** |
| **B) ASD traits** | **Estimates** | **Lower 95% CI** | **Upper 95% CI** | **p** |
| Average Causal Mediation Effects | 8.13 | 6.14 | 10.44 | **<.001** |
| Average Direct Effects | 4.14 | 1.16 | 6.95 | **.005** |
| Total Effects | 12.27 | 10.44 | 14.06 | **<.001** |
| Prop.Mediated | .66 | .48 | .90 | **<.001** |
| **C) Anxiety symptom counts** | **Estimates** | **Lower 95% CI** | **Upper 95% CI** | **p** |
| Average Causal Mediation Effects | 1.84 | .35 | 3.45 | **.012** |
| Average Direct Effects | .81 | -1.24 | 2.79 | .428 |
| Total Effects | 2.65 | 1.42 | 3.89 | **<.001** |
| Prop.Mediated | .69 | .12 | 1.71 | **.012** |
| **D) ODD symptom counts** | **Estimates** | **Lower 95% CI** | **Upper 95% CI** | **p** |
| Average Causal Mediation Effects | .27 | -.26 | .80 | .304 |
| Average Direct Effects | .91 | .16 | 1.66 | **.017** |
| Total Effects | 1.18 | .70 | 1.64 | **<.001** |
| Prop.Mediated | .23 | -.22 | .82 | .304 |

Sensitivity analysis indicated that while the detected mediating effect of coordination ability on ASD traits was robust to unmeasured confounding variables (ρ>.49), the detected mediating effects on ADHD symptoms and anxiety symptoms was sensitive to unmeasured confounders, with small correlations (ρ>.24 for both the ADHD and anxiety models) between unmeasured confounding variables and outcome variables (ADHD or anxiety symptoms) being sufficient to invalidate the detected mediating effect.

Mediation analysis also revealed that coordination ability was a partial mediator of FSIQ, PIQ and VIQ scores, mediating 40% of the effect of having an ND-CNV on FSIQ, and PIQ, and 38% of the effect on VIQ (Table 4).

**Table 4.** Results of mediation analysis on the effect of having an ND-CNV on A) Full Scale IQ B) Performance IQ, C) Verbal IQ with coordination ability as a mediator. Average Causal Mediation Effects corresponds to the indirect effect through coordination ability, while Average Direct effects corresponds to the direct effect of having an ND-CNV on the psychopathology variable of interest.

| **A) FSIQ** | **Estimates** | **Lower 95% CI** | **Upper 95% CI** | **p** |
| --- | --- | --- | --- | --- |
| Average Causal Mediation Effects | -7.12 | -12.17 | -2.58 | **.002** |
| Average Direct Effects | -10.86 | -17.06 | -4.81 | **<.001** |
| Total Effects | -17.98 | -22.02 | -13.98 | **<.001** |
| Prop.Mediated | .40 | .14 | .70 | **.002** |
| **B) PIQ** | **Estimates** | **Lower 95% CI** | **Upper 95% CI** | **p** |
| Average Causal Mediation Effects | -6.68 | -11.79 | -2.08 | **.005** |
| Average Direct Effects | -9.89 | -16.03 | -3.39 | **.005** |
| Total Effects | -16.57 | -20.76 | -12.39 | **<.001** |
| Prop.Mediated | .40 | .12 | .76 | **.005** |
| **C) VIQ** | **Estimates** | **Lower 95% CI** | **Upper 95% CI** | **p** |
| Average Causal Mediation Effects | -6.21 | -11.14 | -1.81 | **.006** |
| Average Direct Effects | -10.28 | -16.66 | -3.75 | **.004** |
| Total Effects | -16.49 | -20.71 | -12.18 | **<.001** |
| Prop.Mediated | .38 | .11 | .74 | **.006** |

However, these analyses were sensitive to confounding with small correlations between unmeasured confounding variables and FSIQ (ρ>.21), VIQ (ρ>.17), and PIQ (ρ>.19) being sufficient to invalidate the detected mediation findings.

Previous studies have found no evidence that IQ is associated with levels of psychopathology in children with an ND-CNV (Niarchou et al., 2014). In order to validate that the mediation analyses were robust to the variable used as a mediator, we constructed a second set of models where FSIQ was included instead of coordination ability as a mediator of the effect of ND-CNV status on psychopathology. In agreement with the previous findings, we found no evidence that FSIQ was a mediator of ADHD (ACME=.25, p=.549), ASD trait (ACME=.44, p=.466), anxiety (ACME=.22, p=.574), or ODD symptoms (ACME=.44, p=.478).

## Discussion

This study shows that difficulties with coordination are common in children with ND-CNVs, with the great majority (91%) of children with an ND-CNV screening positive for coordination problems. We also present evidence that coordination ability is associated with increased ADHD and ASD traits, and lower FSIQ, PIQ and VIQ scores. Coordination difficulties were elevated across all ND-CNV genotypes and genotype or CNV type (deletion or duplication) was not a significant predictor of coordination ability. Importantly, we present evidence that coordination ability is a partial mediator of the effect of ADHD symptoms and ASD traits, along with FSIQ, PIQ and VIQ, and a full mediator of anxiety symptoms, while we found no evidence for mediation of ODD symptoms.

The high rates of coordination difficulties across genotypes, and lack of specificity of ND-CNV genotype on coordination abilty, indicates that neuromotor deficits are a common outcome across ND-CNVs. We found no evidence for an effect of gender on coordination ability, which differs from studies in the general population where DCD is more commonly seen in boys (Missiuna et al., 2008; Tsiotra et al., 2006). It is important to note that rates of premature birth were also high in the ND-CNV group at 54%, Prematurity has been linked to delays in motor development, and coordination difficulties later in life (Edwards et al., 2011; Goyen & Lui, 2009), but we found no effect of prematurity on either screening positive for coordination problems or overall coordination ability. Additionally, while 19% of controls screened positive for coordination problems, this is within the range of prevalence estimates for developmental coordination disorder in the general population, which has been reported as raging from 2%-20%, depending on criteria used (Blank et al., 2019).

Considering links with psychopathology, children with a ND-CNV showed elevated ADHD, ASD trait, anxiety and ODD symptoms compared to controls. Within the ND-CNV group, higher numbers of ADHD, and ASD symptoms were associated with greater motor coordination difficulties, but this was not the case for anxiety or ODD. These results are similar to research in non-genotype selective samples, where high rates of coordination difficulties have been observed in children with ADHD (Watemberg, Waiserberg, Zuk, & Lerman-Sagie, 2007) and/or ASD (Sumner et al., 2016), but differ from findings in children with 22q11.2 deletion, where an association with anxiety was found (Cunningham et al., 2017).

Additionally, coordination difficulties were found to be a partial mediator of the effect of ND-CNV status on ADHD symptoms and ASD traits and a full mediator of anxiety symptoms. No evidence for mediation was found for ODD symptoms. These results may suggest that coordination difficulties are intrinsically linked to the development of ADHD, ASD and anxiety symptoms, but not ODD symptoms, at least in individuals with a ND-CNV.

However it is important to note that the identified mediating effects on ADHD and anxiety symptoms may need to be viewed with caution, as relatively low correlations between any unmeasured (and therefore not accounted for in the mediation models) confounding variable and either ADHD or anxiety symptoms would be sufficient to invalidate the findings.

Importantly, there was no evidence found for mediation of the effect of having an ND-CNV on psychopathology by FSIQ. The lack of a mediating effect of FSIQ on psychopathology also agrees with previous work in 22q11.2 deletion, where it was found that FSIQ was not associated with levels of psychopathology (Niarchou et al., 2014). This helps us validate that our mediation analysis is not prone to detecting false positive mediating effects.

Poor coordination was also found to be associated with FSIQ, VIQ and PIQ in the ND-CNV group. This agrees with previous studies of children with coordination difficulties, not selected due to genotype, and with previous research in 22q11.2DS (Cunningham et al., 2017; Roizen et al., 2011), where associations between FSIQ and motor performance have been found. The presented results, and previous research in 22q11.2DS, support the idea that within ID populations, level of intellectual impairment is associated with poorer motor coordination (Vuijk, Hartman, Scherder, & Visscher, 2010). It may be of note that we found significant associations between coordination ability and raw scores on the matrix reasoning task, while no relationships were found between coordination ability and the block design, vocabulary or similarities tasks. Mediation analysis also found that coordination ability was a partial mediator of the effect of ND-CNV status on FSIQ, VIQ and PIQ. This agrees with theories that suggest that motor skills are required for the development of higher order cognitive skills, such as mathematical ability (Giles et al., 2018), which would be accounted for in PIQ, or language development (Rowe et al., 2008) as accounted for by VIQ.

A number of different interpretations of our mediation findings are possible. First, aberrant development of motor coordination skills as a consequence of a ND-CNV may have cascading impacts on later development of other skills, like cognition, attention and social functioning. Second, motor coordination difficulties trigger environmental risk factors, such as social exclusion and bullying, which may increase risk of psychopathology. Thirdly, both motor coordination problems and impairment in IQ and psychopathology may be the result of the same underlying genetic cause (pleiotropy), although the manifestation of the symptoms may occur at different stages during development. Such pleiotropic effects probably impact on brain development. For example, the cerebellum has been shown to be important in many motor as well as non-motor functions (Miall, Reckess, & Imamizu, 2001; Stoodley, 2016) and, therefore, damage to this region has cross domain effects. It is also possible that multiple processes account for the findings, possibly at different developmental stages and in different ways for each trait. It is for example noteworthy that we did not find evidence of mediation of motor coordination problems for ODD.

We are not able to further distinguish between these hypotheses in the current study, and the cross-sectional design does not allow for insights into order of appearance of different impairments, but delays in motor development are often observed from very early in development. It would be fruitful to conduct a randomised control trial where an intervention for motor coordination difficulties is delivered at an early age to a high-risk group of children with ND-CNVs and changes in psychopathology and cognitive function are established.

This is the only study to investigate the relationships between coordination difficulties and psychopathology and IQ in children with CNV’s that confer risk for psychiatric disorders. The presence of sibling controls for comparison, and detailed assessment of psychopathology are additional strengths. However, the DCDQ is a measure of overall coordination that is completed by a parent, and it therefore cannot allow for direct insights to be gained into underlying sensorimotor deficits that are present in the individual children. To address this, further research should investigate the quality of movement (kinematics), using detailed assessment of fundamental motor control processes. An additional limitation was that the control group was significantly older than the ND-CNV group. However, this should have had limited effect on screening positive or negative for coordination problems, as parents are always asked to rate children with respect to their peers, and screening is based on age dependant thresholds. Younger children require lower scores to screen positive than older children, allowing for a degree of internal control for improvements in ability due to age. In addition, age was included as a covariate in all analyses. Finally, it was not possible to conduct full neurological assessments on the children. Therefore, we cannot rule out other contributing physical or neurological problems that would affect coordination.

In summary, we found very high rates of coordination difficulties in children with CNVs that are associated with risk of neurodevelopmental disorder and which constitute a significant portion of the caseload of clinical geneticists. These problems were elevated across all ND-CNV genotypes and associated with risk of IQ impairment and psychopathology. Furthermore, the association between ND-CNVs and anxiety symptoms was fully mediated by coordination ability, while the association between having an ND-CNV and lower FSIQ, PIQ, and VIQ scores, along with ADHD symptoms and ASD traits was partially mediated by coordination ability. The immediate clinical implication of these findings should be increased vigilance for motor impairments in children with ND-CNVs that are associated with mental health disorders, so that appropriate support can be introduced as early as possible. It may be that motor interventions could help the development of cognitive skills and reduce risk for the development of psychopathology.

## Acknowledgements

We are very grateful for all the families that took part in the IMAGINE-ID study. We would like to thank Dr Samuel Chawner for valuable input and help. We would also like to thank the Core Services team at the MRC Centre for Neuropsychiatric Genetics and Genomics. The work was carried out in collaboration with the National Centre for Mental Health, a collaboration between Cardiff, Swansea and Bangor Universities and funded by Health & Care Research Wales (Welsh Government).

