## Supplementary Table 1 for "Coordination difficulties, IQ and psychopathology in children with high-risk Copy Number Variants"

Supplementary Table 1. Rates of indicated DCD in each CNV group.

| *Genotype* | *DCD* | | ***Total*** |
| --- | --- | --- | --- |
|  | Indicated DCD | No DCD |  |
| 15q11 BP2-BP3 duplication | 0 0 % | 1 100 % | 1 |
| 15q11.2 deletion | 19 90.5 % | 2 9.5 % | 21 |
| 15q11.2 duplication | 3 100 % | 0 0 % | 3 |
| 15q13.3 deletion | 11 91.7 % | 1 8.3 % | 12 |
| 15q13.3 duplication | 7 77.8 % | 2 22.2 % | 9 |
| 16p11.2 deletion | 24 92.3 % | 2 7.7 % | 26 |
| 16p11.2 distal deletion | 7 100 % | 0 0 % | 7 |
| 16p11.2 duplication | 13 100 % | 0 0 % | 13 |
| 1q21.1 deletion | 10 100 % | 0 0 % | 10 |
| 1q21.1 duplication | 12 85.7 % | 2 14.3 % | 14 |
| 22q11.2 deletion | 10 83.3 % | 2 16.7 % | 12 |
| 22q11.2 distal deletion | 3 100 % | 0 0 % | 3 |
| 22q11.2 duplication | 13 86.7 % | 2 13.3 % | 15 |
| Kleefstra (9q.34.3 deletion) | 4 100 % | 0 0 % | 4 |
| NRXN1 (2p16.3 deletion) | 8 88.9 % | 1 11.1 % | 9 |
| TAR deletion | 1 100 % | 0 0 % | 1 |
| TAR duplication | 9 100 % | 0 0 % | 9 |
| ***Total*** | 154 91.1 % | 15 8.9 % | 169 |
