## Supplementary Table 2 for "Coordination difficulties, IQ and psychopathology in children with high-risk Copy Number Variants"

Supplementary Table 2. Regression results for DCDQ score predicted by A) Full Scale IQ (FSIQ), B) Performance IQ (PIQ), C) Verbal IQ, D) Block design raw score, E) Matrix reasoning raw score, F) Similarities raw score G) Vocabulary raw score, with age as a covariate.

| **A)** | | **DCDQ Score** | |
| --- | --- | --- | --- |
| *Predictors* | *ß* | *95% CI* | *p* |
| Age | .23 | .08 – .39 | **.004** |
| FSIQ | .21 | .05 – .36 | **.011** |
| Observations | | 157 | |
| R^2^ / adjusted R^2^ | | .074 / .062 | |

| **B)** | | **DCDQ Score** | |
| --- | --- | --- | --- |
| *Predictors* | *ß* | *95% CI* | *p* |
| Age | .24 | .08 – .40 | **.004** |
| PIQ | .20 | .04 – .36 | **.015** |
| Observations | | 157 | |
| R^2^ / adjusted R^2^ | | .070 / .058 | |

| **C)** | | **DCDQ Score** | |
| --- | --- | --- | --- |
| *Predictors* | *ß* | *95% CI* | *p* |
| Age | .21 | .06 – .37 | **.007** |
| VIQ | .17 | .01 – .32 | **.036** |
| Observations | | 158 | |
| R^2^ / adjusted R^2^ | | .063 / .051 | |

| **D)** | | **DCDQ Score** | |
| --- | --- | --- | --- |
| *Predictors* | *ß* | *95% CI* | *p* |
| Age | .14 | -.03 – .31 | .101 |
| Block design | .15 | -.02 – .32 | .077 |
| Observations | | 155 | |
| R^2^ / adjusted R^2^ | | .058 / .046 | |

| **E)** | | **DCDQ Score** | |
| --- | --- | --- | --- |
| *Predictors* | *ß* | *95% CI* | *p* |
| Age | .12 | -.05 – .29 | .161 |
| Matrix reasoning | .18 | .01 – .35 | **.041** |
| Observations | | 155 | |
| R^2^ / adjusted R^2^ | | .065 / .052 | |

| **F)** | | **DCDQ Score** | |
| --- | --- | --- | --- |
| *Predictors* | *ß* | *95% CI* | *p* |
| Age | .12 | -.05 – .29 | .177 |
| Similarities | .17 | -.00 – .35 | .053 |
| Observations | | 156 | |
| R^2^ / adjusted R^2^ | | .064 / .052 | |

| **G)** | | **DCDQ Score** | |
| --- | --- | --- | --- |
| *Predictors* | *ß* | *95% CI* | *p* |
| Age | .15 | -.02 – .32 | .087 |
| Vocabulary | .13 | -.04 – .30 | .124 |
| Observations | | 156 | |
| R^2^ / adjusted R^2^ | | .056 / .043 | |
